# Infra-slow brain–heart–gut electrophysiological interactions reveal a coordinated multisystem physiological network in humans

**DOI:** 10.64898/2026.04.15.718683

**Authors:** Giovanni Sitti, Laura Pitti, Diego Candia-Rivera

## Abstract

Growing evidence indicates that brain continuously interacts with other physiological systems through neural and non-neural pathways. The brain–heart and brain–gut axes play a central role in homeostasis, allostasis and behaviour, but also in cognitive aspects including emotion and decision-making. Disruptions in these axes have been linked to a wide range of cardiovascular, neurological, and psychiatric disorders. Despite this evidence, triadic crosstalk between the brain, heart, and gut remains largely unexplored. Brain activity, cardiac autonomic fluctuations, and gastric rhythms all exhibit slow temporal components in resting state, suggesting that brain–heart–gut electrophysiological interactions may occur over timescales from the infra-slow (0.01-0.1 Hz) physiological range.

Using non-invasive electrophysiological recordings from 28 healthy participants at rest, we extracted time-varying power dynamics describing the activity of the three organs: brain alpha power, cardiac sympathetic and parasympathetic indices, and the power of the gastric rhythm. Statistical associations among these organs were quantified using the maximal information coefficient across the extended temporal delay range. Physiological interactions were confirmed using surrogate-based testing, which allowed us to construct the network topology of interactions between the three organs. Our findings show that triadic brain–heart–gut interactions form a multi-directional network at infra-slow timescales, shaping resting state activity. This study offers one of the first insights into the physiology of brain–heart–gut interplay, providing a methodological baseline for the development of more comprehensive biomarkers based on network dynamics capable of linking pathological conditions to dysregulation across multiple organ systems.

**Abstract figure legend:** 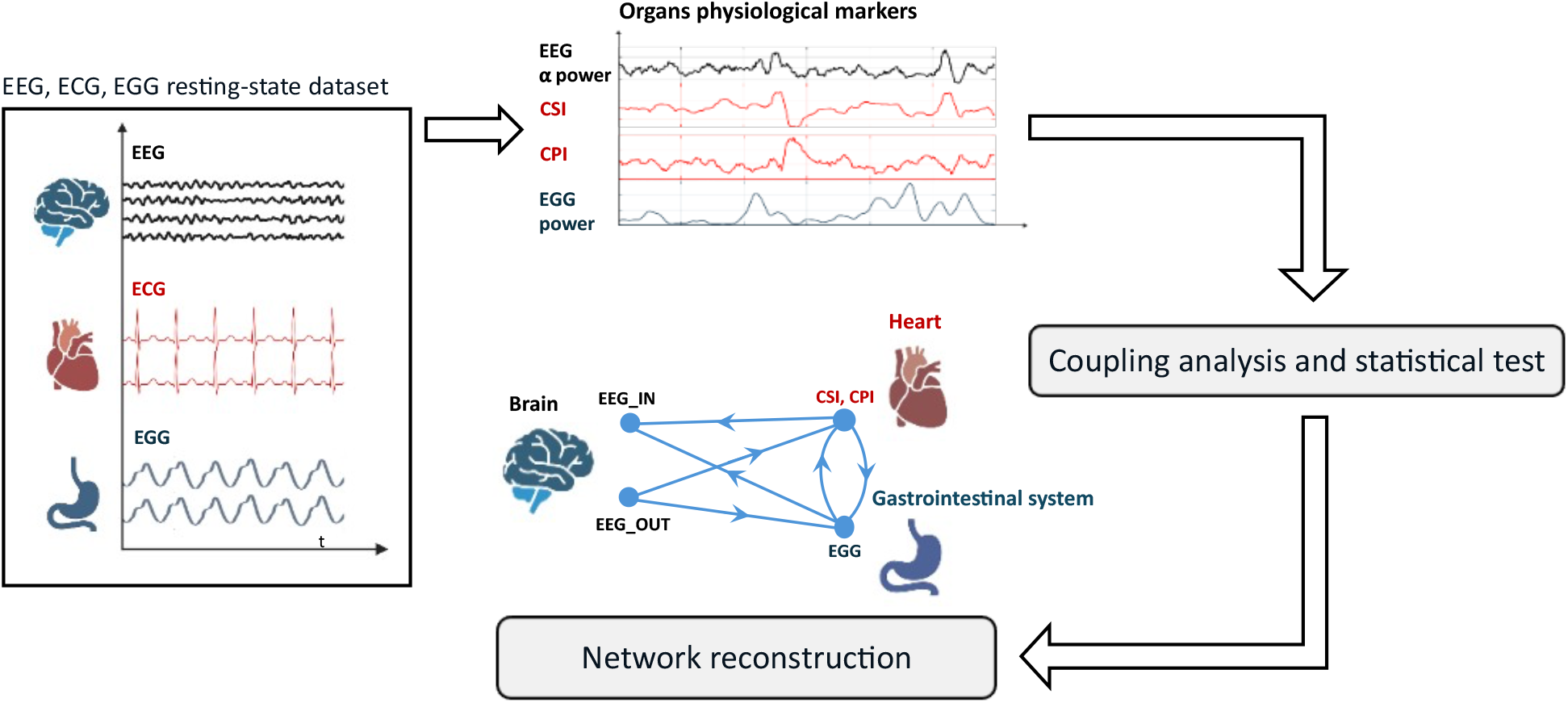

Simultaneous resting-state electroencephalographic (EEG), electrocardiographic (ECG), and electrogastrographic (EGG) recordings were processed to extract time-resolved physiological markers for each organ: EEG alpha-band power for the brain, cardiac sympathetic and parasympathetic indices (CSI, CPI) for the heart, and EGG power for the gastrointestinal tract. Coupling between time series was then quantified, and statistical significance was assessed using a surrogate-based method. Significant couplings were subsequently integrated to construct a large-scale network representation, summarizing the strength, temporal delays, and directionality of the predominant electrophysiological interactions among the three organs.

**Key points summary:**

- First in-human, non-invasive investigation of parallel brain–heart–gut electrophysiological interactions in awake, healthy individuals.
- We analysed simultaneous electroencephalographic (EEG), electrocardiographic (ECG) and electrogastrographic (EGG) recordings and quantified strength and temporal scale of the derived time-series associations, to construct a large-scale network of interactions.
- We found that brain–heart–gut interactions extend into the infra-slow (0.01 - 0.1 Hz) range, indicating that spontaneous fluctuations in the electrophysiological activity of one organ at rest are typically followed by corresponding changes in the other two.
- We found a consistent brain–heart–gut network topology across participants, with multidirectional interactions and bodily dynamics converging toward midline central-posterior brain regions.
- These findings provide one of the first endeavours in understanding the physiology of brain–heart–gut interactions, and a methodology with strong biomarker development potential.

## Introduction

In network neuroscience, neural activity is interpreted as the result of large-scale network dynamics operating across multiple spatial and temporal scales (Bassett & Sporns, 2017). However, this field has been predominantly confined to the brain, despite its function constantly influence and its influenced by other organs activity through neural, autonomic, hormonal, and visceral pathways (Candia-Rivera *et al*., 2024; Subramanian *et al*., 2025). The human organism can therefore be seen as a network of interacting physiological systems, whose coordinated dynamics are essential for maintaining homeostasis and shaping cognitive and behavioural states (Blanken *et al*., 2021). In particular, the brain–heart and brain–gut axes play a central role in regulating emotional processing, cognition, and overall physiological functions (Azzalini *et al*., 2019). Growing evidence shows that the two axes share common neural pathways and comorbidities. Indeed, both the heart and the gastrointestinal tract can communicate with the brain mainly through the sympathetic and parasympathetic branches of the autonomic nervous system, via both partially overlapping afferent and efferent pathways (Müller *et al*., 2022; Haruki & Ogawa, 2023). Peripheral signals are conveyed through the spinal cord to brainstem nuclei, including the nucleus tractus solitarius and the dorsal motor nucleus of the vagus, and are subsequently integrated within central autonomic network regions such as the thalamus, hippocampus, amygdala, and insular cortex (Chen *et al*., 2021). Moreover, mental health has been linked to gastrointestinal and cardiovascular manifestations (Fang & Zhang, 2024; Banellis *et al*., 2025), while neurodegenerative disorders have also been linked to heart failure and gut dysbiosis (Kim *et al*., 2023; Munoz-Pinto *et al*., 2024). Heart rate variability (HRV) alterations have been associated to gastrointestinal disorders (Sadowski *et al*., 2021) and cerebrovascular diseases linked to gut dysbiosis (Wei *et al*., 2023).

Some quantitative evidence of brain–heart–gut interactions exist, such as the modulation of coupling strength between the three organs across sleep stages, based on bowel sounds (Wang *et al*., 2025), the identification of brain–heart–gut metabolic networks in mild cognitive impairment (Li *et al*., 2026), the interplay between stomach pH, HRV and emotions (Porciello *et al*., 2024), the parallel report of HRV and the phase of the gastric rhythm on fMRI (Rebollo *et al*., 2018), and the association between HRV and brain-gastric phase coupling (Rebollo & Tallon-Baudry, 2022). Despite these preliminary findings, the electrophysiology of brain–heart–gut interaction remains to be characterized, underscoring the need for a robust quantitative framework to investigate the spatial and temporal scales of this triadic interplay.

We hypothesize that brain–heart–gut interactions can occur at slow (0.1-1 Hz) and infra-slow (0.01-0.1 Hz) physiological timescales in resting-state humans. Indeed, infra-slow activity has been correlated to blood-oxygenation-level-dependent signals and with resting-state network dynamics in fMRI, and modulate cortical excitability and alpha band resting state activity (Picchioni et al., 2011; Hiltunen et al., 2014; Van Putten et al., 2015). Similarly, cardiac autonomic fluctuations and gastric slow waves exhibit dynamics at slow and infra-slow temporal scales (Wolpert et al., 2020). Moreover, hormonal signalling such as the hypothalamic–pituitary–adrenal (HPA) axis generates ultradian rhythms through delayed feedback mechanisms, further contributing to slow and infra-slow cross-system modulation (Lightman & Conway-Campbell, 2010).

The aim of this study is therefore to characterize temporal structure, strength, and directionality of brain–heart–gut interactions, adopting a multimodal non-invasive approach based simultaneous electroencephalographic (EEG), electrocardiographic (ECG), and electrogastrographic (EGG) resting-state human recordings. Particularly, the gastrointestinal system is assessed through EGG, which reflects intrinsic gastric slow-wave activity generated by interstitial cells of Cajal, typically occurring around 0.05 Hz, modulated by enteric nervous system activity and smooth intestinal muscle movements (Wolpert *et al*., 2020). A dedicated preprocessing pipeline was developed to extract time-varying power dynamics from the brain as measured from EEG, from the gut measured from EGG, and from the heart, exploiting cardiac autonomic measures derived from the ECG.

In this study, we used Maximal Information Coefficient (MIC), since it can detect both linear and nonlinear associations without assuming specific model structures (Reshef *et al*., 2011; Cao *et al*., 2021). Using MIC, we tested time-delayed coupling within a temporal window of 120 seconds, containing slow and infra-slow temporal scales. Spatial and temporal information obtained has been exploited to construct a network representation describing the functional circuitry connecting the brain, heart, and gut. This study provides a new step toward a more comprehensive understanding of brain–heart–gut interactions within a unified framework, introducing a reproducible approach for quantifying multi-organ interplay across temporal scales. This framework may support future investigations aimed at identifying network level biomarkers, improving the characterization of physiological and pathological states.

## Methods

### Ethical approval

Prior to data collection, ethics approval was obtained from relevant ethics committee at Anglia Ruskin University (approval code: FST/FREP/17/75). Experimental procedures were developed in accordance with the Declaration of Helsinki.

### Dataset description

The data has been collected as part of an existing study (Todd *et al*., 2021), and are publicly accessible on Figshare website as “Body image and implicit interoception dataset “ (Todd & Aspell, 2020). The participants included in the original dataset comprised 36 right-handed individuals (15 men and 21 women, ranged in age from 19 to 40 years). None of the individuals reported a history of neurological or psychiatric disorders, or present use of medications that can affect diet or weight, and no physical disorders, as assessed through a screening questionnaire administered prior to participation in the original study. The data used consists of EEG, EGG and ECG human recordings, EEG activity has been recorded with 32 active scalp electrodes (FP1, FP2, F7, F3, FZ, F4, F8, FC5, FC1, FC2, FC6, T7, C3, CZ, C4, T8, TP9, CP5, CP1, CP2, CP6, TP10, P7, P3, PZ, P4, P8, PO9, O1, OZ, O2, PO10) positioned according to the international 10-20 system. EGG and ECG signals were recorded simultaneously with EEG. For the EGG, six disposable cutaneous electrodes were placed over the abdominal surface with a six-lead configuration. ECG activity was recorded using three disposable cutaneous electrodes arranged in a standard three-lead configuration. The ECG and EGG recordings shared a common ground electrode. For each subject, recordings were stored in BrainVision format and consisted of 38 channels at a sampling rate of 500 Hz: 3 EGG channels, 3 ECG channels, 32 EEG channels. Physiological data were recorded for approximately 12 minutes, where participants were instructed to avoid voluntary movements and to keep their eyes open (except for natural blinking). They were encouraged to let their mind wander freely and to refrain from engaging in structured mental activities.

The dataset used in this study comprised recordings from 28 participants, since a subset has been excluded due to missing data, specifically incomplete EEG, ECG and/or EGG recordings.

### EEG processing

All preprocessing procedures were implemented using MATLAB-R2024a. The preprocessing pipelines for EEG, ECG, and EGG signals were developed separately and were primarily based on functions available in FieldTrip open-source MATLAB toolbox (Oostenveld *et al*., 2011). EEG signals have been band-pass filtered between 1 and 45 Hz using a fourth order Butterworth filter. Powerline interference was suppressed using a notch filter at 50 Hz. Physiological artifacts removal was performed using Independent Component Analysis (ICA), with the extended infomax algorithm (Jung *et al*., 1998). Eye blinking and cardiac artifacts were identified and removed through visual inspection of the independent components, considering their temporal dynamics, spectral content, and spatial topographies. To further attenuate residual high transient noise, a wavelet-denoising procedure in the ICA domain was applied (Gabard-Durnam *et al*., 2018). Each selected independent component time series was decomposed using a stationary wavelet transform. The removed signal portion was then projected back to sensor space using the ICA mixing matrix, allowing subtraction of the reconstructed artifacts from the EEG data. Bad channel removal was then performed in a semi-automatic manner, estimating channel correlation respect to each neighbour and by visual inspection. Channels identified as bad were removed and subsequently reconstructed using spline-based spatial interpolation. The clean data was average-referenced and the final pre-processed EEG data has been saved for subsequent analyses (Candia-Rivera *et al*., 2021).

Spectral power was estimated for all EEG channels using a frequency resolution of 0.5 Hz. A Hanning taper was applied to each sliding time window to reduce spectral leakage. The time window length was fixed at 2 s with 50% overlap. The power time series for each EEG channel were obtained by the integration of the EEG power spectrum across the α frequency range (8–12 Hz).

### EGG processing

EGG signals were pre-processed in MATLAB exploiting Fieldtrip open-source toolbox “EGG_Scripts” (*Wolpert* et al., *2020*). Signals were visually inspected to identify potential artificial artifacts, or electrode-related noise. A preliminary check of spectral content has been performed by computing the Fast Fourier Transform of each channel, to verify that the dominant spectral peak of each channel fell within the expected normogastric frequency range (0.033 to 0.066 Hz) (*Richter* et al., *2017*; *Rebollo* et al., *2018*). Some channels that did not show a dominant spectral peak within the expected normogastric frequency range were discarded (*Wolpert* et al., *2020*). This ensured that only channels reflecting reliable gastric myoelectrical activity were retained for subsequent processing and analysis. Subsequently, signals were band-pass filtered at 0.03 - 0.07 Hz, to isolate the information content in the normogastric range, which characterizes the gastric myoelectrical activity, eliminating respiratory (∼0.3 Hz), cardiac rhythms (∼1.5 Hz) and higher frequency related noise. To characterize the temporal fluctuations of gastric activity, time-frequency analysis was performed on the pre-processed EGG signals using FieldTrip, using a sliding-window multi-taper convolution approach with an Hanning taper and next power of two padding. A window length of 40 seconds was used, providing sufficient spectral resolution for slow gastric rhythms. Spectral power was integrated across the frequency band of interest (0.03 – 0.07) Hz, yielding a single time-varying measure of gastric power. This produced the EGG power, representing slow fluctuations of gastric rhythmic activity over time. The mean EGG power envelope of the remaining channels for each subject has been computed and used for the coupling analysis.

### ECG processing

R-peak locations in the recordings were obtained using an ECG automatic peak detection algorithm (R-DECO toolbox) (Moeyersons *et al*., 2019), where R-peak identification is performed using an envelope-based algorithm that flattens the ECG signal and enhances QRS complexes, improving robustness to noise. The sequence of RR intervals was then computed as the difference between successive R-peak instants, and IBI inter-beat-interval (IBI) series were constructed based on the R-to-R-peak durations. First order Poincaré plot - representing IBI value against the subsequent interval has been computed (IBI_*i*_ vs. IBI_*i+1*_, where *i* is the index of each of the IBIs identified in the ECG). Three time-varying features have been extracted from the Poincaré plot: the cardiac cycle duration, measured as the distance to the origin (CCD), and the minor (SD_1_) and major (SD_2_) ratios of the ellipse that quantify the short- and long-term fluctuations of HRV, respectively.

The time-varying fluctuations of the distance to the origin and the ellipse ratios were computed with a sliding-time window T = 15s, as follows:

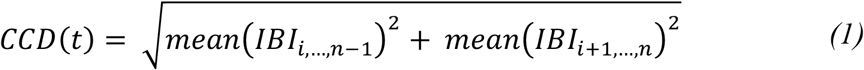

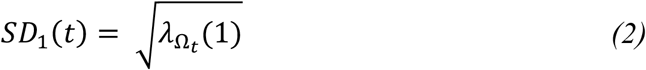

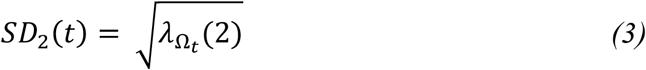

where 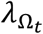 is the matrix with the eigenvalues of the covariance matrix of *IBI*_{*i*,…,*n*–1}_ and *IBI*_{*i*+1,…,*n*}_, with Ω_*t*_: *t* − *T* ≤ *t*_*i*_ ≤ *t*, and n is the length of IBI in the time window Ω_t_.

In this study, the ‘exact’ approach has been used according to the implementation described in (Candia-Rivera et al., 2025b), which computes the standard covariance matrix giving the covariance between each pair of elements. The distance to the origin CCD_0_ and ellipse ratios SD_01_ and SD_02_ corresponds to the computation on the whole recording and were computed to re-centre the time-resolved estimations of CCD, SD_1_ and SD_2_. Then, the CPI and the CSI are computed as follows:

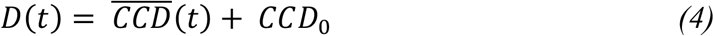

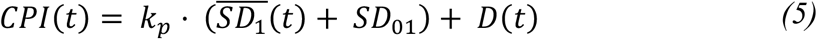

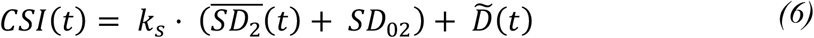

where 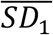 and 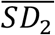 are the demeaned *SD1* and *SD2* and 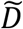 is the flipped D with respect to the mean. The coefficients *k*_*p*_=10 and *k*_*s*_ = 1 determine the contribution of fast and slow HRV oscillations respect to changes in baseline cardiac cycle duration (CCD) (Candia-Rivera et al., 2025b). CPI represents the minor axis of the ellipsoid, with its variability reflecting the fast fluctuations in heart rate variability associated with parasympathetic tone. CSI represents the major axis of the ellipsoid, with its variability reflecting the slow fluctuations in heart rate variability associated with sympathetic tone.

### Lagged coupling analysis

Prior to coupling analysis, EEG alpha power was smoothed using a 15 sec moving average to suppress faster oscillations and match the temporal resolution of cardiac indices. All signals were then resampled to 1 Hz using spline interpolation. The median group power spectral density of all timeseries has been finally computed to examinate the spectral content and thus understand the temporal scale of the single organ electrophysiological activity.

Maximal Information Coefficient (MIC), a non-parametric measure of statistical association, has been used for coupling analysis since it is able to detect both linear and nonlinear dependencies without assuming a predefined model between signals. This makes it well suited for a physiological framework, where interactions are often nonlinear and complex (Reshef *et al*., 2011). MIC explores all possible grid partitions of two variables within a finite dataset to search for the maximal mutual information, which is then normalized. Specifically, for a pair of variables (*X, Y*), the MIC of *X* and *Y* is defined as:

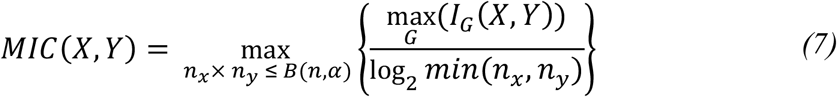

where *n*_*x*_ and *n*_y_ are the number of bins on the x-axis and y-axis, respectively. *G* represents a *n*_*x*_ × *n*_y_ grid on (*X,Y*), *I*_*G*_(*X, Y*) denotes the mutual information under the grid *G*, and *B*(*n, α*) is a function of data size *n* and is equal to *n*^*α*^, (0 <*α* <1), which limits the maximum number of bins. The normalization term ensures that *MIC* lies in the range (0-1). MIC has been implemented using minepy matlab toolbox (Reshef *et al*., 2011). The MIC computation has been done setting *α* =0.6, and the grid search refinement factor c = 15, based on default parameter settings (Reshef *et al*., 2011). To account for physiological delays, coupling was computed over a lag window of ±120 sec, and the sign of the optimal lag defines the interaction direction. In addition, as a control measure, we confined the search for an optimal delay in the range 0-10s and 100-120s. For each EEG channel, couplings were computed between the alpha-band power and each autonomic or gastric index (CSI, CPI, and EGG power). For brain-body interactions (e.g., EEG–CSI, EEG–CPI, EEG–EGG), a single maximum was selected across the delay range, allowing the sign of the optimal lag to reveal the direction of interaction (brain → body for positive delays, body → brain for negative delays). In addition, coupling between cardiac autonomic indices and gastric activity was computed, assuming bidirectional interactions, by independently identifying the optimal lag in both the positive and negative delay ranges (Gut → Heart for positive delays, Heart → Gut for negative delays). Couplings were computed for each channel. Based on the minimum number of participants showing significant coupling, a data-driven threshold was defined to identify significant results at the group level.

### Network construction

Physiological time-series have been modelled as nodes in directed weighted graphs. Since CSI and CPI derive from the same cardiac signal, they cannot be treated as independent interacting systems; thus, no direct links were considered between them. An edge from node *i* to node *j* was included only if the coupling between the corresponding signals was statistically significant, and the sign of the corresponding optimal lag gives the direction of the edge *i* → *j*. The weight C_ij_ of each edge was defined as the value of the coupling computed at the optimal lag_ij_. An example of large scale “Brain - Body Network” built according to these rules is shown in Figure 1.

**Figure 1:**
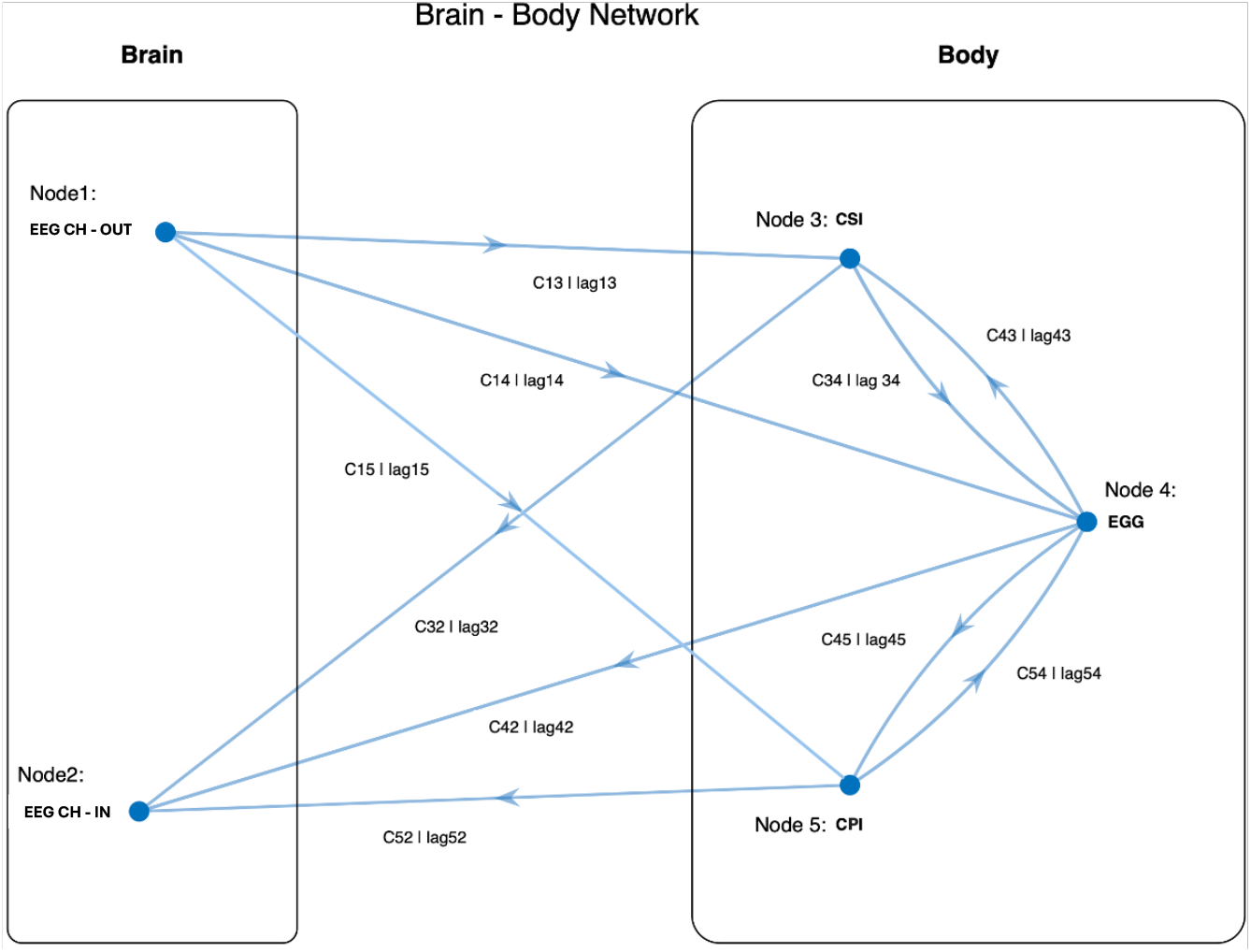
Schematic representation of a brain–body network derived from lagged coupling analyses in this study, showing all possible connections permitted by the model. To enhance clarity and interpretability, one input (EEG CH-IN) and one output (EEG CH-OUT) node were included for the brain system.

### Statistical analysis

For each subject and for each pair of physiological signals a surrogate based statistical test was performed. Surrogate versions of the original signals at the best lag were generated using the Iterative Amplitude Adjusted Fourier Transform (IAAFT) algorithm (Sparacino *et al*., 2025). For each surrogate, the coupling value was therefore calculated, yielding to surrogate distribution representing the null hypothesis, against which the real coupling value was statistically evaluated using a Monte Carlo approach to reject the presence of spurious correlations, avoiding false positive. An empirical one-sided p-value was estimated as the proportion of surrogate values exceeding the real coupling value. The coupling has been considered statistically significant when *p* < *α*, where an *α* = 0.05 as been fixed (Candia-Rivera & Valenza, 2022).

### Data availability statement

The data used in this study is part of an open access dataset, accessible on Figshare as “Body image and implicit interoception dataset” (Todd & Aspell, 2020) https://aru.figshare.com/articles/dataset/Body_image_and_implicit_interoception_dataset/13296314/2. A detailed description of the dataset and protocol can be gathered from the original study (Todd *et al*., 2021),

## Results

Figure 2 displays the time series obtained after the processing procedure. CSI and CPI are obtained from ECG, EEG alpha power from EEG, and the EGG power derived from the gastric signal. The spectral content of alpha power peaks around 0.01 Hz, whereas bodily signals exhibit peak frequencies below 0.01 Hz (Figure 3), indicating that signals are dominated by slow temporal variations over tens of seconds, compatible with infra-slow physiological fluctuations (<0.1 Hz).

**Figure 2:**
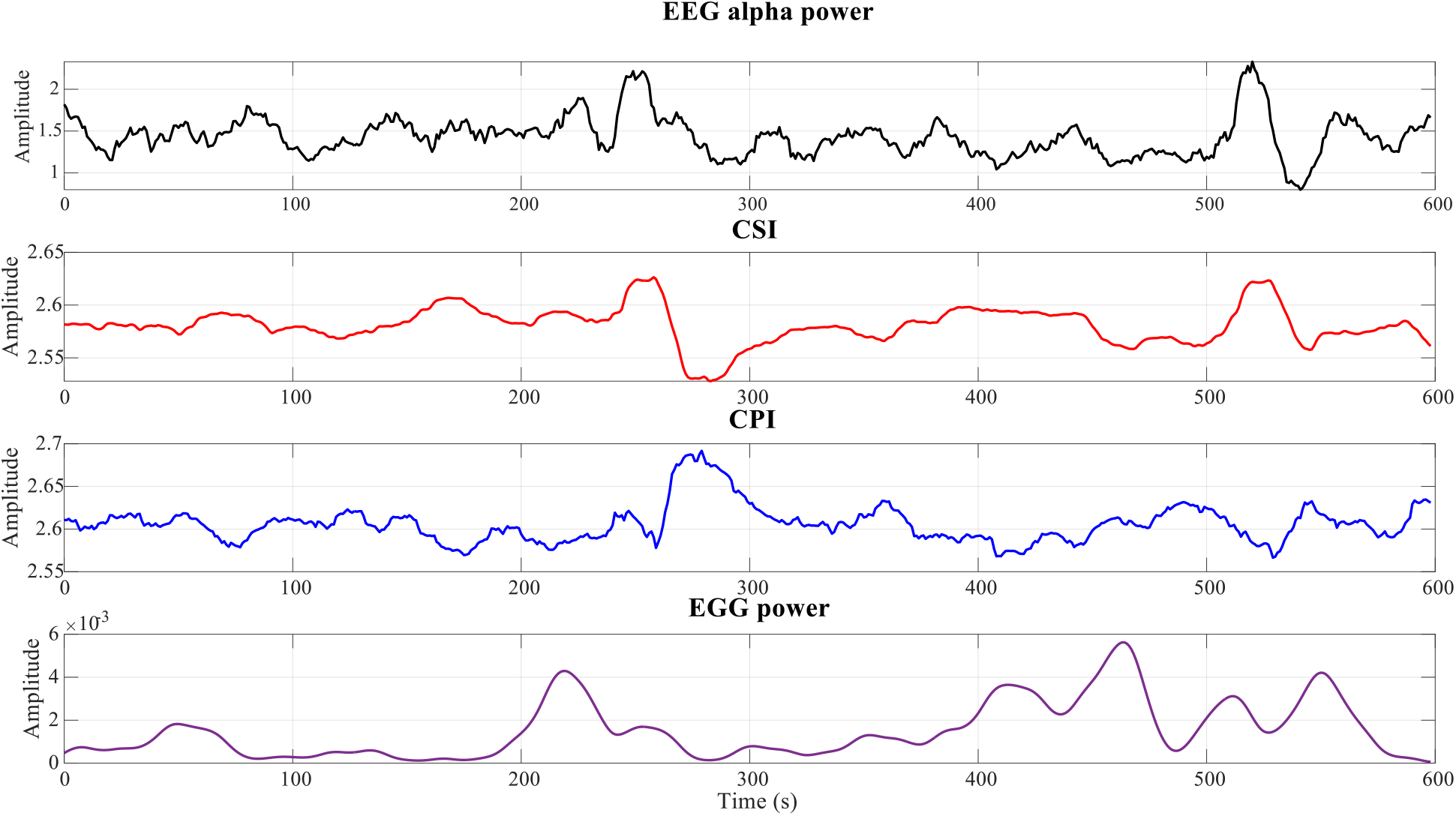
Single subject time series resulting from the preprocessing pipeline: EEG alpha power, representing brain resting state activity, CSI and CPI, representing heart activity, and the EGG power, representing the gastrointestinal system electrophysiological activity.

**Figure 3:**
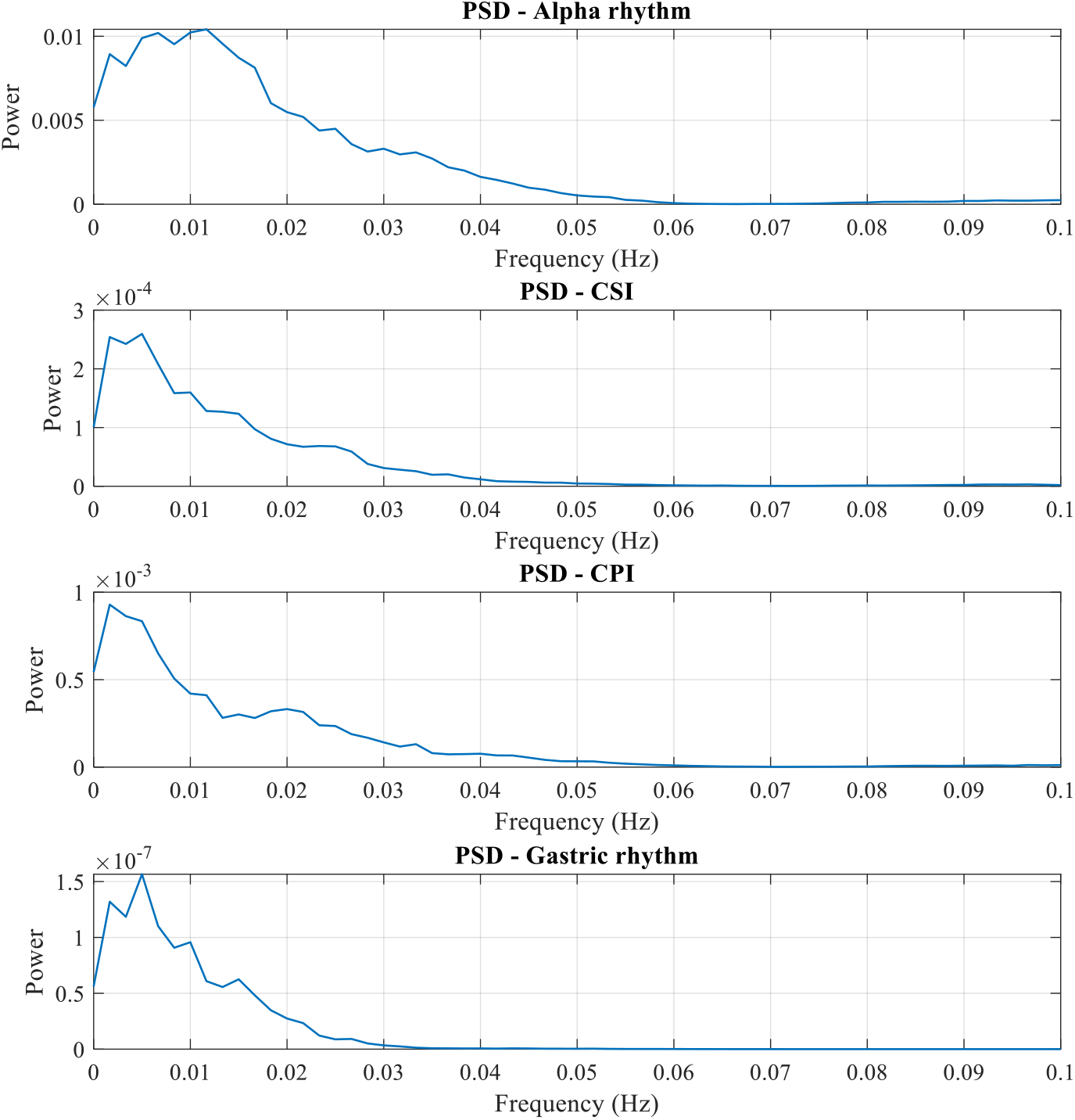
Group median power spectral density of the EEG alpha power, cardiac sympathetic and parasympathetic indices (CSI and CPI) and the EGG power.

For each dyadic coupling with the brain, the scalp topographic maps in Figure 4 show the number of participants having statistically significant coupling with each peripheral physiological signal per channel. This allowed us to assess how consistently significant couplings appeared across subjects on each channel.

**Figure 4:**
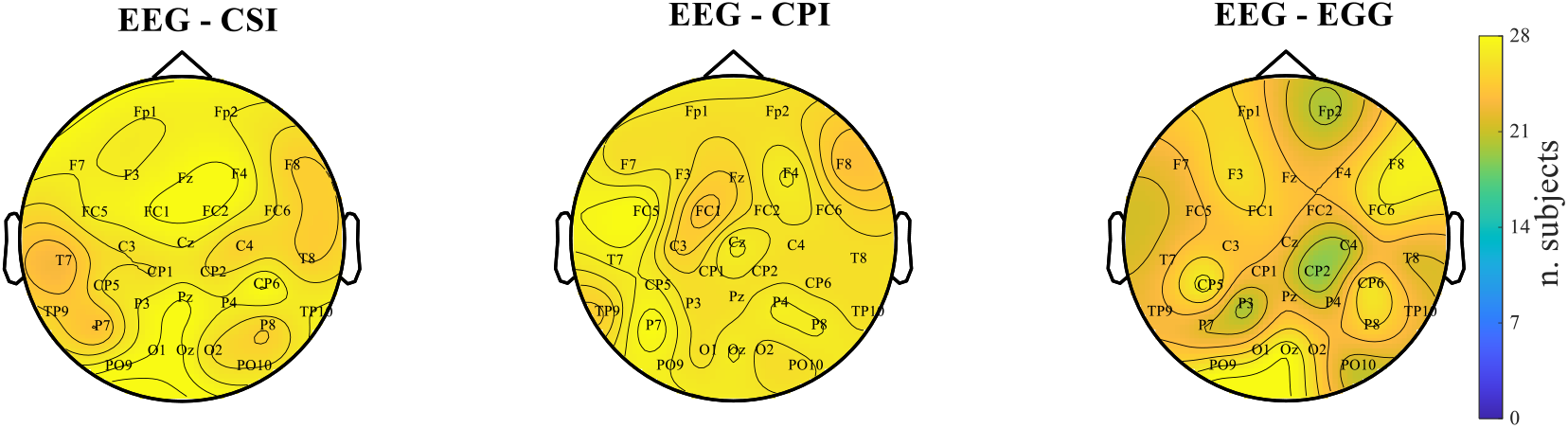
Alpha band topographic maps showing the number of participants having statistically significant coupling with each peripheral physiological signal per channel.

These results show that, for each channel, there are at least 19 out of 28 significant subjects (approximately 70% of the sample). Based on this observation, a 70% threshold was adopted for the group-level analysis, meaning that only EEG channels that showed statistically significant coupling in at least 70% of participants were included. The corresponding median group topographic maps are presented in Figure 5, showing the median maximum coupling value for each channel in the alpha band for each type of coupling and sign of delay, allowing visualization of the spatial distribution of brain→body and body→brain interactions.

**Figure 5:**
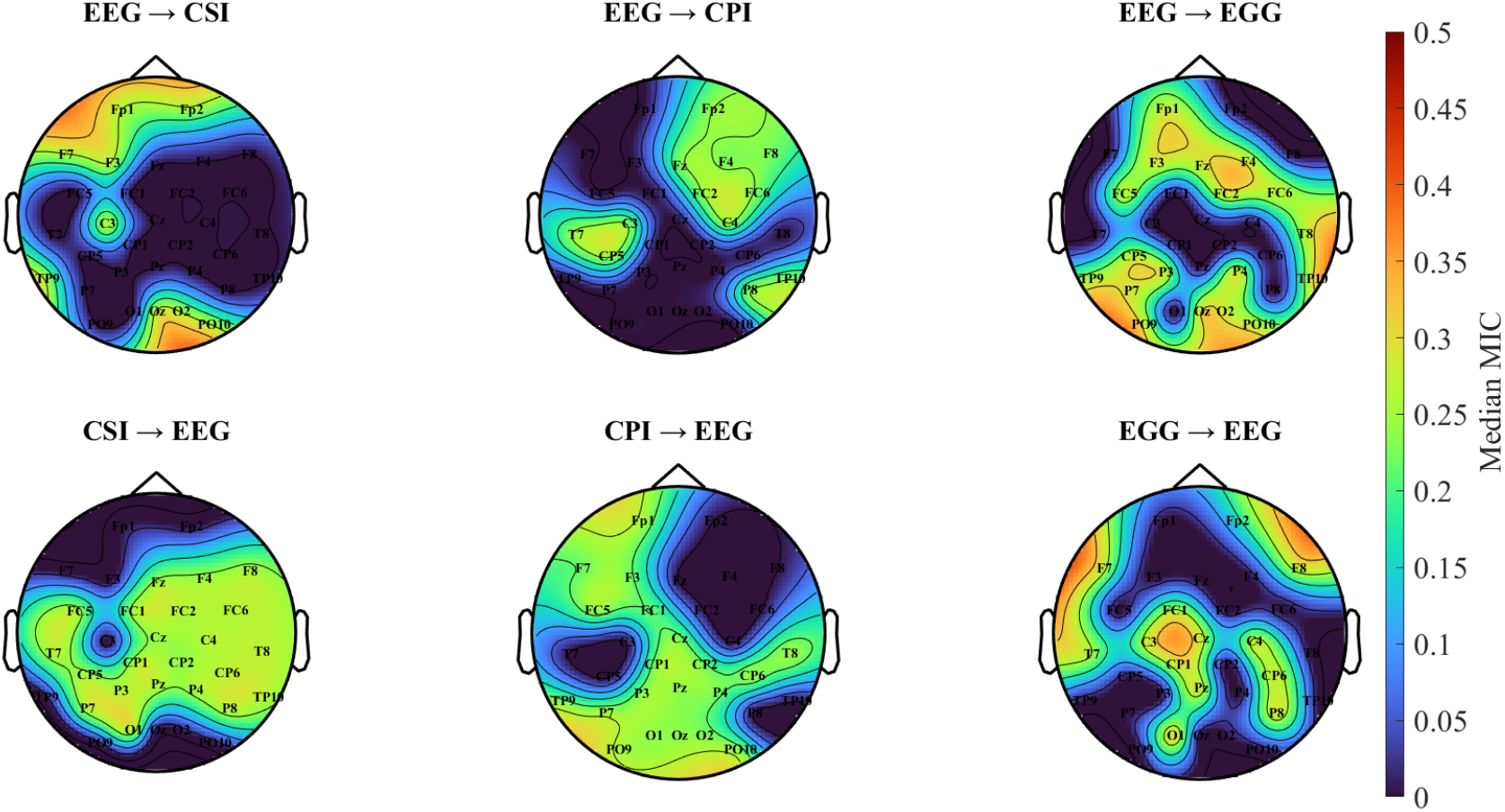
Group median topographic maps showing the maximum information coefficient per EEG channel, for each direction with respect to the cardiac sympathetic and parasympathetic indices (CSI and CPI) and the gastric rhythm (EGG).

EEG–EGG coupling shows structured spatial patterns, with central clusters (C3, FC1, Cz, Pz), left temporo-parietal (F7, T7), right frontal (Fp2, F8), and posterior regions (PO9, TP9, P7, Oz, O2, PO10), as well as a frontal cluster (F3, Fp1, Fz, F4, FC2). CPI→EEG showed more distributed central and frontal scalp regions, while EEG→CPI right frontal organization. CSI→EEG shows distributed central areas, and EEG→CSI stronger parietal–occipital (Oz, O2, PO10) and frontal areas (Fp1, Fp2, F7, F3).

To investigate which scalp regions might be involved in an information flow driven by both the heart and the gut, only the EEG channels that were in common across the corresponding coupling maps are shown in Figure 6. We found a midline central-occipital cluster (FC1, Cz, CP1, Pz, O1) where bodily signals converge.

**Figure 6:**
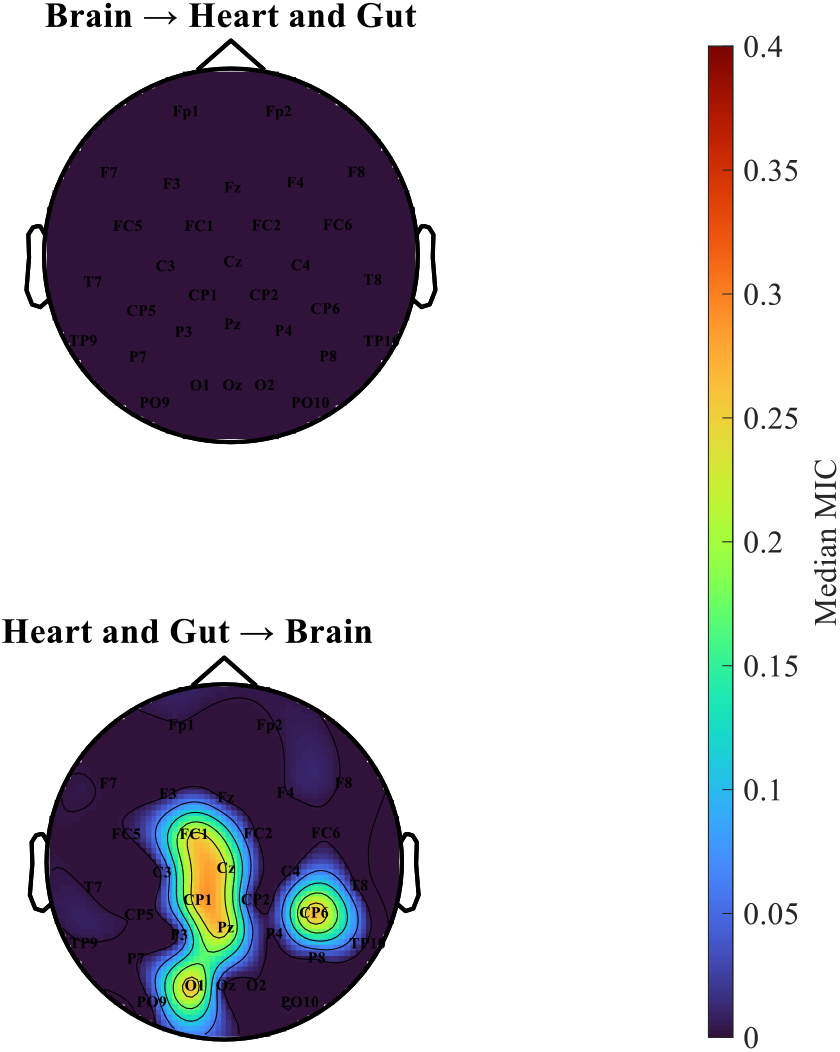
Group median topographic maps displaying convergent scalp regions coupled with the heart and gut, as quantified by EEG – CSI, EEG – CPI, and EEG – EGG couplings.

Histograms in Figure 7 show the number of participants exhibiting maximal coupling at specific delays for each coupling type. For brain → body and body → brain couplings, only the EEG channel with the maximum coupling was considered for simplicity. This representation allows visualizing the temporal structure and directionality of brain-body and heart-gut interactions across subjects. The distribution of the percentage of participants showing significant coupling at defined timescales is presented in Table 1.

**Figure 7:**
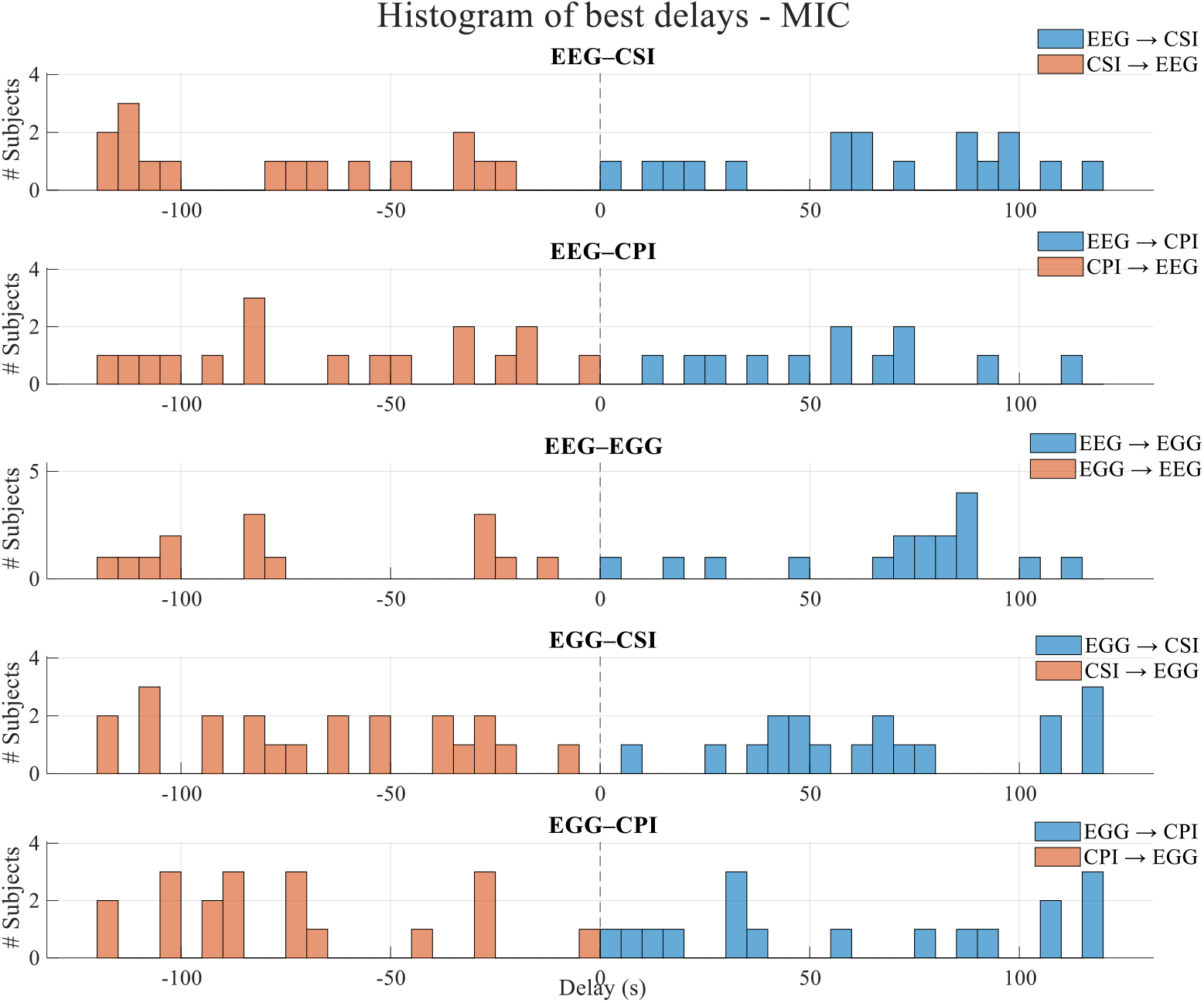
Histograms of the predominant delays showing the number of subjects at each specific lag value, for each coupling type.

**Table 1:**
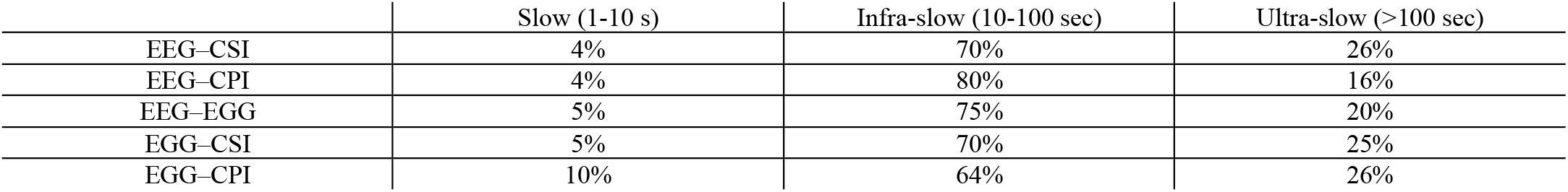
Distribution of the percentage of participants of coupling delays across multiple timescales.

|  | Slow (1-10 s) | Infra-slow (10-100 sec) | Ultra-slow (>100 sec) |
| --- | --- | --- | --- |
| EEG-CSI | 4% | 70% | 26% |
| EEG-CPI | 4% | 80% | 16% |
| EEG-EGG | 5% | 75% | 20% |
| EGG-CSI | 5% | 70% | 25% |
| EGG-CPI | 10% | 64% | 26% |

Figure 8 shows the group median network representation of the electrophysiological large-scale coupling within the brain–heart–gut system, summarizing how information flows across systems at group level (see Supplementary Material to visualize individual network topologies). Brain is modelled as a simple 2 nodes system. An output node, represented by the EEG channel having the strongest median maximum coupling in the positive-delay direction (Oz), and an input node, as the EEG channel having the strongest median maximum coupling in the negative delay direction (O1). Cardiac system is represented by two nodes corresponding to the autonomic indices CSI and CPI. Gastric activity is represented by the EGG node.

**Figure 8:**
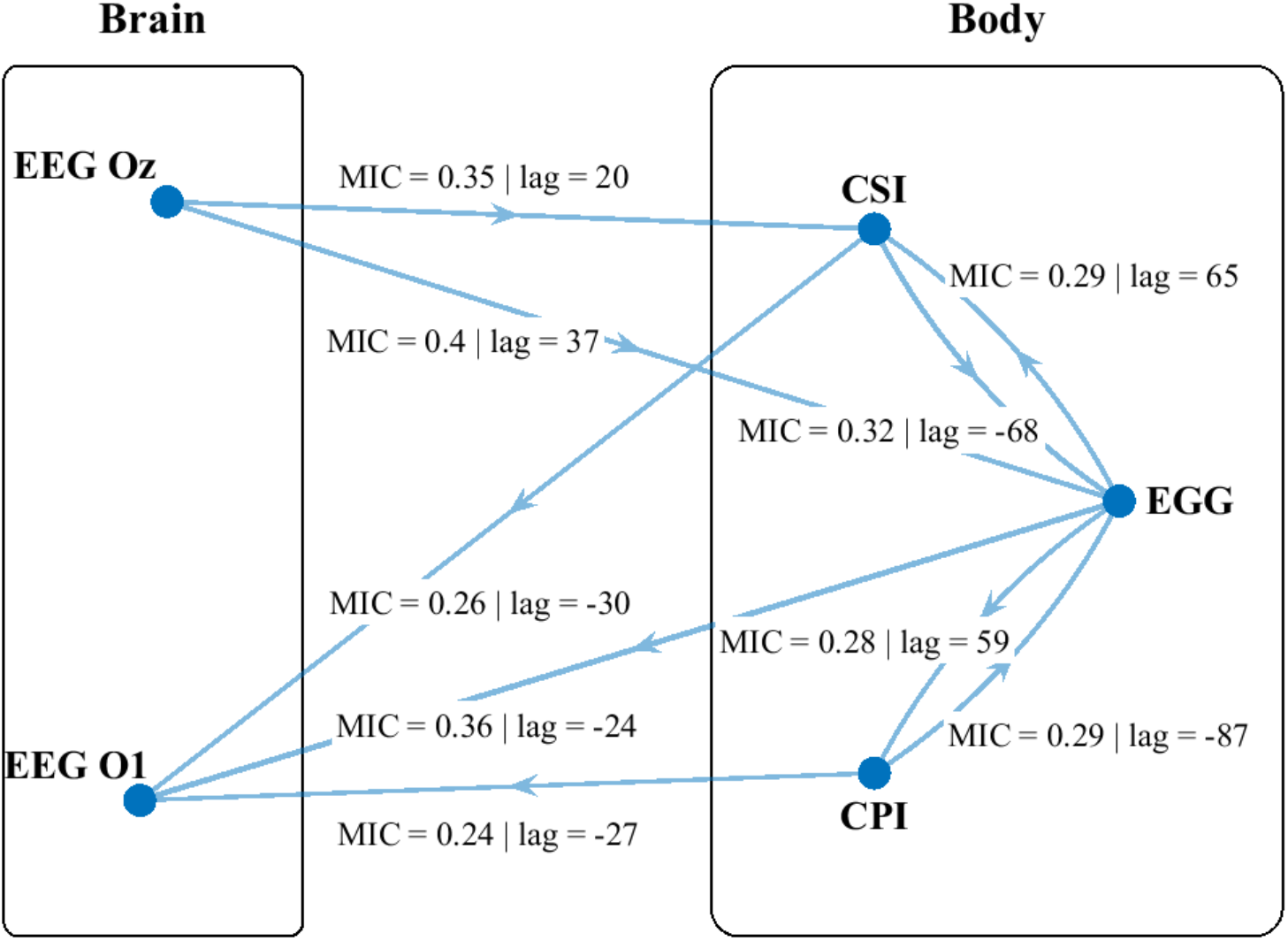
Group median Brain – Heart – Gut network topology, highlighting the strength, direction, and characteristic delays of significant information exchange among the cortical, cardiac, and gastric systems. The network reveals a distributed multidirectional organization, where posterior cortical regions (Oz, O1) interact with both autonomic cardiac indices (CSI, CPI) and gastric rhythm (EGG). The reported delays, spanning from 20 to 87 seconds, indicate that couplings occur predominantly at the infra-slow time scale.

## Discussion

Robust interactions between cardiac, gastric, and brain activity were observed at infra-slow timescales, revealing a coherent and spatially structured pattern of coupling during rest. These findings support the hypothesis that cardiac and gastric dynamics are intrinsically linked to brain activity and potentially contribute to the organization of resting-state physiological networks. Physiological couplings were consistently detected across participants, with statistically significant interactions observed across all EEG channels, albeit at different timescales. This consistency indicates that brain–heart–gut interactions constitute a reproducible and spatially organized physiological phenomenon. Moreover, significant clusters of interactions emerged in both the body→brain and brain→body directions, suggesting the presence of a large-scale, bidirectional organization linking these three systems.

We found a topographic cluster in the ascending direction from the heart and gut to the brain, converging to a midline central-posterior cluster, which may indicate the existence of a common monitoring architecture of visceral activity or to shared pathways through which visceral informations are jointly processed (Engelen *et al*., 2023). Previous studies showed a clear association of certain brain regions to gastric and cardiac dynamics, including the posterior cortex, such as the posterior portion of the cingulate cortex and the praecuneus, but also the somatosensory cortex (Ladabaum *et al*., 2001; Wang *et al*., 2008; Park *et al*., 2014; Cao *et al*., 2019, 2022; Al *et al*., 2021). This convergence may reflect the brain’s role in monitoring bodily signals to maintain physiological regulation and support cognitive, emotional and behavioural states. Such an organization is consistent with theoretical frameworks of interoception, in which distributed cortical regions integrate visceral inputs to support predictive regulation of bodily states and self-related processes (Craig, 2009; Barrett & Simmons, 2015). For example, in the case of bodily modulations triggering nutrient intake (Hadjieconomou *et al*., 2020; Ly *et al*., 2023; Lyu *et al*., 2024; Woodie *et al*., 2024), or the coordinated patterns of gastric activity, feelings of disgust or fear, and increased HRV illustrate this dynamic brain–body interplay (Porciello *et al*., 2024).

Physiological interactions between the brain and peripheral organs were characterized by infra-slow latencies spanning from a few seconds to over 100 seconds, with substantial inter-individual variability. Notably, both positive and negative optimal delays were observed across all coupling types, indicating the absence of a dominant directionality while preserving a consistent spatial organization of interactions. The infra-slow range emerged as the predominant timescale of coupling, accounting for 60–80% of observed delays and exhibiting median latencies between approximately 20 and 90 seconds. These results point to a dynamic equilibrium in which central and peripheral systems continuously and reciprocally modulate each other through slow bidirectional exchanges of physiological information.

Such infra-slow dynamics are consistent with known properties of large-scale brain activity, where slow fluctuations are thought to regulate neural excitability and shape perceptual sensitivity (Monto *et al*., 2008). In parallel, slow EEG dynamics have been linked to vascular, hemodynamic, and metabolic processes (Obrig *et al*., 2000; Palva & Palva, 2012; Nikulin *et al*., 2014), suggesting that the observed delays may reflect integrated neurophysiological and systemic mechanisms. The timescales identified here also align with prior reports of delayed interactions across bodily systems, including ∼20-second lags in cardiovascular regulation (Porta *et al*., 2011; Czarnek *et al*., 2021), 20–60 second delays linking peripheral afferents to cardiac responses (Candia-Rivera *et al*., 2025*a*), and respiration-driven brain modulations unfolding across slow and infra-slow ranges (Birn *et al*., 2008; Candia-Rivera *et al*., 2022).

Importantly, longer delays exceeding 100 seconds may not solely reflect direct neural communication but could also arise from slower biochemical signalling pathways. Endocrine processes, such as pulsatile cortisol secretion governed by the hypothalamic–pituitary–adrenal axis, operate through delayed feedback loops and can influence both brain and body over tens of seconds to minutes (Lightman & Conway-Campbell, 2010). More broadly, hormonal signalling spans multiple temporal scales, from rapid pulsatile events to ultradian rhythms, providing a plausible substrate for the extended latencies observed (Lamont & Amir, 2017). Additionally, some delays may reflect harmonics of underlying slower oscillatory coupling, whereby fundamental interactions at shorter periods give rise to secondary peaks at integer multiples. Together, our findings could also be interpreted as a potential effect of non-electrophysiological dynamics, meaning that infra-slow brain–body interactions may emerge from the joint interplay of neural, autonomic, and endocrine mechanisms, collectively shaping a temporally extended and multiscale coordination between central and peripheral physiology.

From a methodological point of view, the quantification of interactions among physiological time-series requires appropriate analytical methods. Traditional approaches such as correlation and spectral coherence have been widely used to assess pairwise interactions, but are limited to detect linear relationships (Candia-Rivera *et al*., 2026). Causal reconstruction methods such as Granger Causality rely on model-based assumptions, while Transfer Entropy requires high-dimensional state-space reconstruction, making it computationally expensive and highly sensitive to data length and noise (Runge, 2018). Within this context, the use of the MIC provides a complementary approach to characterize both linear and nonlinear associations without imposing a specific model structure.

Some limitations should be considered in this work. First, all measurements were obtained using non-invasive electrophysiological recordings, thus several relevant subsystems may operate as hidden nodes that influence the detected interactions. These include neural structures not accessible through scalp brain recordings, such as subcortical regions, as well as components of the enteric nervous system that cannot be fully resolved. Second, although temporal delays were used to infer directionality, the present approach does not establish causal interactions in a strict mechanistic sense. Third, we did not perform an exhaustive search of all physiologically plausible couplings across different latencies, given the substantial computational demands such an analysis would entail. Instead, we focused on identifying the strongest coupling within a defined timescale and then validated whether these couplings remained the strongest after surrogate testing. This approach ensures statistical robustness and reduces the likelihood of reporting false positives. Another limitation is that the study focused exclusively on resting-state conditions. Still, the present work represents one of the first systematic attempts to characterize electrophysiological coupling among these three organ systems simultaneously. Future studies could move beyond scalp-level analyses and investigate brain–heart–gut interplay at the whole brain level (Rebollo *et al*., 2018), in order to better identify the specific brain regions involved. Beyond pairwise analyses, higher-order interaction analysis could provide a more complete representation, allowing the characterization of collective dependencies among three or more signals (Rosas *et al*., 2019; Santoro *et al*., 2023). Future research could repeat this analysis in different physiological or task-related conditions to investigate how brain–heart–gut coupling is modulated across experimental states. Network-based measures may therefore support the identification of multiorgan physiological signatures that better reflect the organism’s global state from a unified perspective.

Future research on the electrophysiological interplay between the brain, heart, and gut should aim to integrate this multiscale framework with emerging domains that capture the biological and behavioural complexity of interoceptive regulation. One particularly promising direction concerns the interaction with the gut microbiota, which has been increasingly implicated in shaping neural, autonomic, and endocrine function through the microbiota–gut–brain axis (Cryan *et al*., 2019; Mayer *et al*., 2022). Combining electrophysiological measures with microbiome profiling could help disentangle how microbial composition and metabolite signalling influence infra-slow brain–body dynamics. In parallel, linking these physiological interactions to subjective and behavioural dimensions, such as body awareness, interoceptive accuracy, and body image (Todd *et al*., 2021), may provide critical insight into how large-scale brain–heart–gut coupling contributes to self-related processes and mental health (Candia-Rivera *et al*., 2024). Alterations in these domains are central to conditions such as anxiety, depression, and eating disorders, where disrupted interoceptive signalling has been repeatedly reported (Khalsa *et al*., 2018).

Another key avenue involves the investigation of clinical populations, particularly in relation to cognitive impairment and neurodegenerative disorders. Dysregulation of autonomic and gastrointestinal function can be observed in ageing and dementias (Cox *et al*., 2026). Characterizing how infra-slow brain–body coupling is altered in these conditions could yield novel biomarkers for early detection and disease monitoring, as well as inform mechanistic models linking systemic physiology to neurodegeneration (Foster *et al*., 2017). More broadly, future studies should explore how these interactions evolve across the lifespan, are modulated by factors such as sleep and circadian rhythms, and respond to interventions including neuromodulation, biofeedback, diet, or pharmacological treatments. Integrating multimodal recordings with computational modelling approaches will be essential to capture the nonlinear and bidirectional nature of these systems. Ultimately, advancing this line of research may provide a unified understanding of how distributed physiological networks support cognition, emotion, and health, opening new avenues for precision medicine and systems-level interventions.

## Conclusions

We demonstrate the presence of coordinated electrophysiological interactions between the brain, heart, and gut at infra-slow timescales using a multimodal, non-invasive approach. These findings support the hypothesis that cardiac and gastric dynamics are intrinsically coupled with brain activity and may contribute to the organization of large-scale neural processes. By introducing a framework that captures nonlinear, delayed, and bidirectional interactions, this work provides a methodological basis for investigating multiorgan physiological coupling in humans. More broadly, our results support the view that physiological regulation emerges from distributed network interactions spanning multiple organ systems, extending the principles of network neuroscience toward an organism-level perspective grounded in integrative physiology. This study establishes a foundation for future investigations of brain–body interactions in both health and disease.

## Acknowledgement

L.P. and G.S. were supported by a travel grant from the University of Bologna (Bando Borse di studio per tesi all’estero, Lauree Magistrali - Dipartimento DEI Cesena).

